# Impact of carbon monoxide on early cardiac development in an avian model

**DOI:** 10.1101/2021.12.22.473783

**Authors:** Filipa Rombo Matias, Ian Groves, Mari Herigstad

## Abstract

Carbon monoxide (CO) is a toxic gas that can be lethal in large doses and may also cause physiological damage in lower doses. Epidemiological studies suggest that CO in lower doses over time may impact on embryo development, in particular cardiac development, however other studies have not observed this association. Here, we exposed chick embryos *in ovo* to CO at three different concentrations (1ppm, 8ppm, 25ppm) plus air control (4 protocols in total) for the first nine days of development, at which point we assessed egg and embryo weight, ankle length, developmental stage, heart weight and ventricular wall thickness. We found that heart weight was reduced for the low and moderate exposures compared to air, and that ventricular wall thickness was increased for the moderate and high exposures compared to air. Ventricular wall thickness was also significantly positively correlated with absolute CO exposures across all protocols. This intervention study thus suggests that CO even at very low levels may have a significant impact on cardiac development.

## INTRODUCTION

Carbon monoxide (CO) is a toxic gas without odour, colour or taste (Penney et al., 2010). While produced endogenously, primarily through heme degradation (Stevenson et al., 2001), exogenous CO exposure in higher doses can be lethal. Haemoglobin (Hb) has a markedly higher affinity for CO than for oxygen, and carboxyhaemoglobin (COHb) is readily formed. Importantly, as CO remains bound and if exposures to the gas persists, COHb continues to rise at the increasing expense of oxyhaemoglobin (OHb) (Penney et al., 2010). CO also shifts the OHb dissociation curve and causes reduced release of oxygen to the tissues (Longo, 1977; Townsend & Maynard, 2002). Together, this can lead to severe hypoxia and death. Toxic mechanisms beyond hypoxia may also contribute to further morbidity, as symptoms can persist or appear even after COHb levels return to normal (S R Thom et al., 1995).

Pregnant women are at an increased risk for CO poisoning due to increased endogenous CO production during gestation (Smollin & Olson, 2008). Acute CO poisoning during pregnancy has been associated with premature delivery and spontaneous abortion, with pregnancy outcome likely dependent on severity of maternal poisoning and foetal age (Smollin & Olson, 2008). Foetal death may occur at nonlethal maternal carbon monoxide exposures (Longo, 1977).

While it is generally acknowledged that CO poisoning can cause severe health consequences or even death, much less is known about lower-level exposures. CO exposures of 6 ppm and lower can impact on vascular function (Bendell et al., 2019) and epidemiological studies report associations between maternal CO exposure and ventricular septal defects in the foetus (Dadvand et al., 2011; Ritz et al., 2002; Zhang et al., 2016) at levels as low as ~1 ppm (Dadvand et al., 2011). Other studies, however, have failed to replicate these findings (Chen et al., 2014). As foetal COHb will, under steady-state conditions, be 10-15% higher than maternal COHb (Longo, 1977), the foetus may be particularly at risk during longer-term exposures.

CO exposure is difficult to study experimentally in humans. The level of CO that is ethical to deliver is limited both in terms of duration and amount. While some work has been done using low-level CO interventions in humans, e.g. (Bendell et al., 2019), such studies would not be feasible in pregnant women. The chick is a common model for developmental research, as the embryo is easily accessible *in ovo* with progressive organ development which is highly conserved with mammals. It is also a good model for CO research, as CO responses in the chick resembles mammalian ones (Stupfel et al., 1982). Also, at Hamburger-Hamilton stage 35 (embryonic day (D)9) the chick embryo heart, with its four chambers, bears a closer structural resemblance to the human heart than other non-mammalian model organisms (Wittig & Munsterberg, 2016). The gaseous environment in which eggs are incubated can be easily controlled, thus further cementing its usefulness as a model for CO research.

The aim of the current study was to interrogate the impact of low-level CO exposure on early development in the chick embryo, particularly focusing on cardiac development.

## METHODS

Fertilized Bovan Brown chicken eggs (Gallus gallus) (Henry Stewart & Co., Norfolk, UK) were used for all experiments. All experiments were performed according to relevant regulatory standards. Eggs were incubated from D0 to D9 of development in an incubator (Thermo Scientific) at 37°C ±0.1°C. The eggs were separated in custom-made air-tight incubation boxes inside the main incubator, each containing a separate 25 mL glass container with ultra-pure water to maintain air humidity. Each box was equipped with two ports, closable with IV 3-way tap valves, used for adjustment of internal gas composition. Throughout the incubation period, each incubation box held either air (control) or CO targeted at the following concentrations: 1ppm, 8ppm and 25ppm. These levels correspond to low CO exposure (on par with UK atmospheric levels), moderate CO exposure (on par with current UK guidelines) and high exposure (half that of common CO home alarm sensor levels (50ppm)). The experiment was run on 4 separate batches of eggs, and the eggs in each batch was divided randomly and equally into the three CO conditions and Air control.

Gas exposure was conducted as follows: CO gas was sampled from a pre-mixed cylinder (1000ppm CO in air, BOC) into a non-pressurized reservoir (2L anaesthetic bag), from which a pre-determined amount was drawn and introduced to the incubators via the ports using a 20mL syringe. The amount of gas required was calculated based on the size of the sealed individual incubators (each box had a volume of 2.7L). To test that target CO levels in the incubators were reached and maintained throughout the experiments, gas levels were measured through one of the incubator ports every 1-2 days of incubation (TPI 716 flue gas analyser, Test Products International Europe Ltd, West Sussex, UK). Every 2-3 days, this was also followed by a venting of the incubators and re-administration of gases to avoid longitudinal drift in oxygen and carbon dioxide content. Also at this point, ultra-pure water levels were assessed and replenished if necessary.

### Measurements

On D9, eggs were removed from the incubator and weighed. Embryos were removed and assessed for viability (presence of heartbeat) immediately prior to termination by decapitation. All nonviable embryos were excluded from further analysis. Viable embryos were harvested, removing any yolk/extraembryonic membranes. The following measurements were obtained prior to histological processing:

#### Weight

Weight was measured using a standard laboratory analytical balance. Full embryo weight was assessed immediately following termination. The heart was then carefully excised, blotted dry and weighed.

#### Staging

Development and morphogenesis were visually assessed using the Hamburger-Hamilton Staging Series (Hamburger & Hamilton, 1992; Martinsen, 2005) as a reference guide. For a subset of embryos, the assigned stage was confirmed by a second researcher, as a quality control.

#### Size

Embryo size was assessed for each viable embryo as a measure of development. Full length was measured from the top of the skull frontal bone to the posterior end of the pelvic bones, toe length (third toe) from the phalangeal-tarsometatarsal joint to the posterior end of the toe, and ankle length from the anterior end of the tarsometatarsus - ankle joint - to the tip of the posterior end of the (third) toe. All measurements were conducted by positioning the embryo and limbs on a grid surface under a stereoscope.

### Histology measurements

#### Histology protocol

Hamburger-Hamilton stage 33-35 hearts were fixed in formalin 10% for 24h at room temperature, then processed for paraffin embedding (see supplement for protocol) in a Leica TP 1020 tissue processor. Tissues were embedded in Leica HistoCore Arcadia H. Haematoxylin and Eosin (H&E) staining were performed manually (see supplement for protocol) on 3μm transversal plane sections obtained by the Leica RM2235 microtome.

#### Ventricular wall thickness

Four hearts were used to optimise the histology procedure and excluded from further analysis. In the remaining hearts, ventricular wall thickness was measured across transverse plane sections from the outer surface to the inner surface of the myocardium at three different points for three different sections using cellSens Dimension (Olympus Life Science, Olympus, Tokyo) through an Olympus IX81 motorized inverted research microscope (Olympus Life Science). Ventricular wall thickness was averaged for each section for statistical comparison. No further measurements of cardiac morphology were taken. Ventricular wall thickness was re-assessed for air control and moderate CO exposure in a separate, subsequent experiment to verify findings.

### Statistical comparisons

Students’ T-tests (one-tailed) were used to compare control and CO exposure levels (Microsoft Excel, Microsoft Corporation, WA, USA). As three comparisons were made for each measurement, corrections for multiple comparisons (Bonferroni method) meant that outcomes needed to meet a threshold of p<0.0167 to be considered significant at an alpha of p<0.05. All p-values are reported as is, but only those meeting the adjusted threshold reported as significant. Correlations between CO level and the end-point measurements of heart weight and ventricular wall thickness were assessed using regression models (SPSS 24.0 (IBM SPSS Statistics for Windows, Armonk, NY, USA).

## RESULTS

Measurements were taken from 44 viable embryos. One embryo was excluded from analysis following an outlier analysis (underdeveloped embryo), leaving 43 embryos for statistical comparison. Four embryos were used to optimise the histology protocol, and a further 4 embryos were excluded from analysis of heart weight and ventricular wall thickness due to failure to dissect complete tissues, leaving 35 embryos for this part of the analysis.

### CO values

For each CO target value, actual CO values deviated with on average +2.4ppm (CO low category, target 1ppm), +1ppm (CO moderate category, target 8ppm) and −6.8ppm (CO high category, target 25ppm). CO values varied slightly between egg batches (see table 1). There was a significant difference in measured CO levels between the low, moderate and high exposures, as expected (one-way ANOVA (F(3,39) = 142.9, p<0.0001), but not between experiments (one-way ANOVA (F(5,37) = 2.088, p=0.089).

**Table 1.** Weight and size measurements (N=43). Means and (SD). Stage is given as median and [25^th^-75^th^ quartile]. Statistical comparisons are between CO and air: *p<0.05, corrected for multiple comparisons, ǂp<0.05, not corrected for multiple comparisons (not significant).

| Measurement | Air (control) | CO, low | CO, moderate | CO, high |
| --- | --- | --- | --- | --- |
| N | 10 | 11 | 11 | 11 |
| CO target (ppm) | 0 | 1 | 8 | 25 |
| CO values (ppm) | 0 (0) | 3.4 (0.9) | 9.0 (3.2) | 18.2 (2.9) |
| Egg weight (g) | 57.33 (3.28) | 58.06 (2.78) | 57.29 (2.40) | 57.45 (2.97) |
| Embryo weight (g) | 1.41 (0.27) | 1.35 (0.23) | 1.27 (0.19) | 1.29 (0.22) |
| Stage (HH) | 35 [35-35] | 34 [33-36] | 33.5 [33-35] | 35 [33-35] |
| Embryo length (mm) | 3.16 (0.34) | 2.96 (0.17)‡ | 2.88 (0.27)‡ | 2.98 (0.24) |
| Ankle length (mm) | 0.67 (0.11) | 0.60 (0.08) | 0.56 (0.08)* | 0.60 (0.07) |
| Toe length (mm) | 0.28 (0.04) | 0.21 (0.11) | 0.23 (0.07) | 0.24 (0.05) |
| Ventricular thickness (µm) | 397 (65) | 460 (44) | 519 (37) | 515 (57) |

### Egg weight, embryo staging and size

We observed no significant difference in egg weight or embryo weight between any of the CO exposures and air control, nor was there any significant difference in stage between any CO exposures and air control. A significant reduction in ankle length (p=0.014) was found for moderate CO compared to air control. Data are presented in Table 1.

### Heart weight and ventricular wall thickness

We observed a significant reduction in heart weight for the moderate CO exposure compared to air control (p=0.013, Figure 1). A significant increase in ventricular wall thickness was found for the moderate (p=0.0005) and high (p=0.005) exposures (Figure 2). This increase in ventricular wall thickness was also found in a subsequent repeat experiment using a moderate CO exposure level (8ppm) compared to air control.

**Figure 1.**
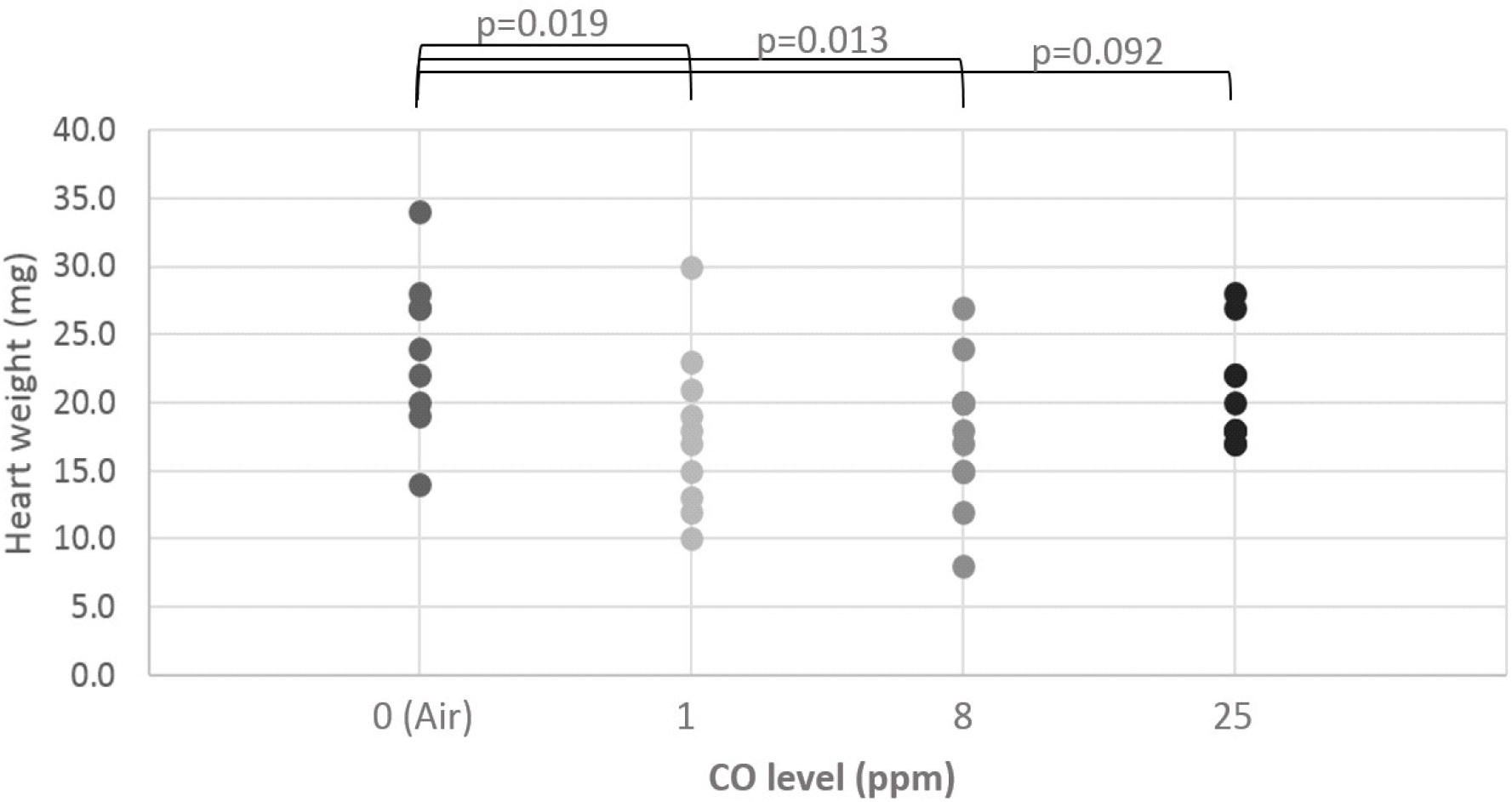
Embryo heart weight (blotted dry, mg). Individual data (N=35) and target CO values (ppm) plus air (control).

**Figure 2.**
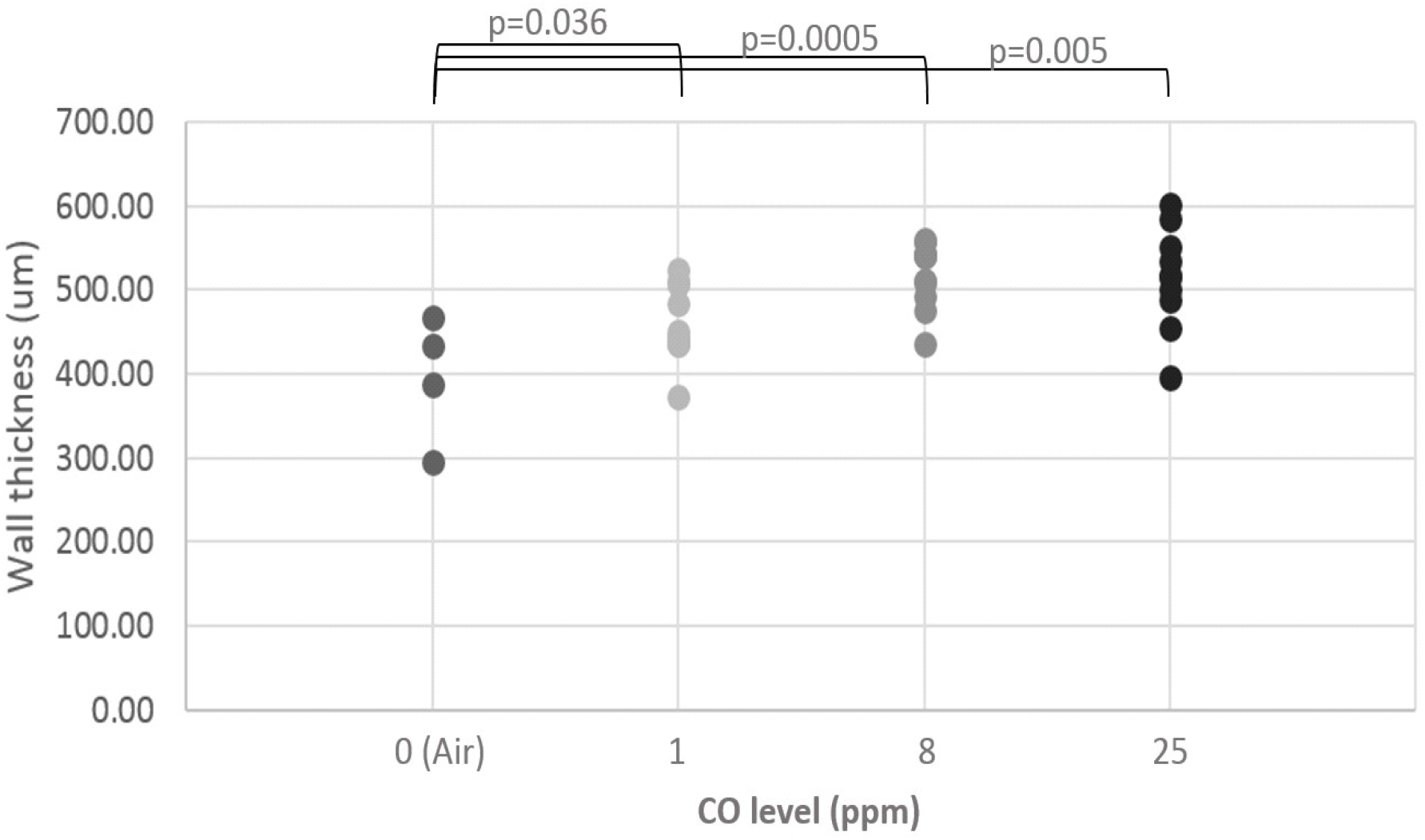
Embryo ventricular wall thickness (μm). Individual data (N=35) and target CO values (ppm) plus air (control).

A significant positive linear correlation was found between the measured CO levels for each batch and ventricular wall thickness (R=0.506, R2=0.256, adjusted R2=0.230, p=0.004, Figure 3), but not for heart weight.

**Figure 3.**
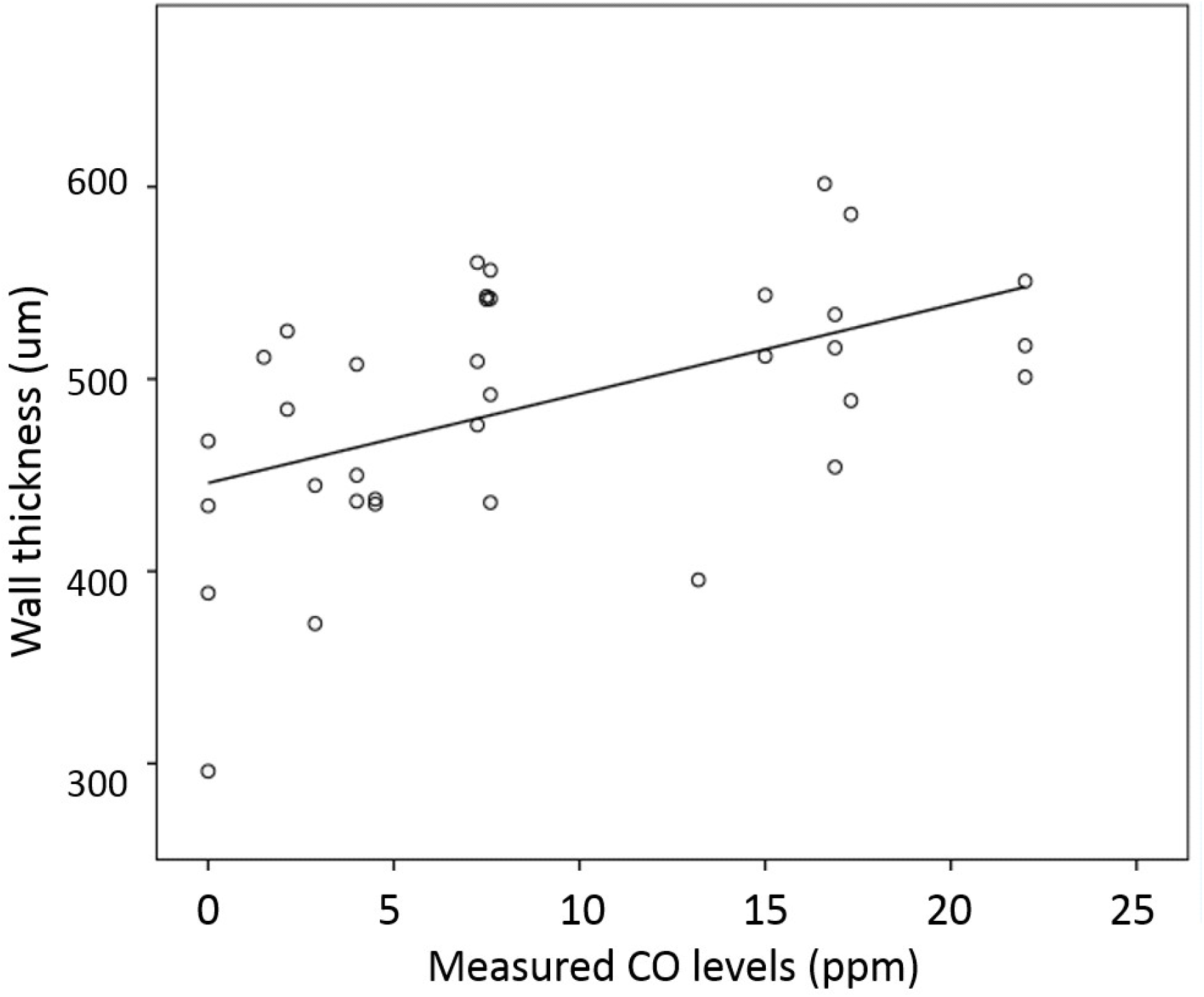
**Ventricular wall thickness (μm) plotted against absolute CO levels** (ppm) for all specimens (N=43). Linear regression line included to show slope of relationship (R2=0.256, p=0.002).

Based on the distribution in Figure 1, we also tested whether there was a U-shaped curvilinear relationship between heart weight and CO level. The trajectory showed a reduction in heart weight in the low and medium CO exposures compared to air control, and almost no difference in heart weight between the highest CO exposure and air control. We did not find a significant relationship between the measured CO levels for each batch (averages across each 10-day incubation period for each batch) and heart weight (R=0.308, R2=0.095, adjusted R2=0.046, p=0.159). However, there was a significant relationship between average CO level across batches and heart weight (R=0.388, R2=0.151, adjusted R2=0.105, p=0.048).

## DISCUSSION

In this study, we show that CO exposure impacts on cardiac development in the chick embryo. We observe that there is a significant and dose-dependent impact on heart weight and structure (ventricular wall thickness), and that ventricular wall thickness correlates with CO exposure.

### Levels of CO

CO has been shown to have an effect on physiology at low concentrations. For example, smokers are exposed to CO through cigarette smoke and typically have persisting elevated levels of CO of 6ppm (exhaled air) or more (Middleton & Morice, 2000). An increase in exhaled CO to this level has been shown to reduce cerebrovascular reactivity and alter fMRI signal in healthy human non-smoking volunteers (Bendell et al., 2019). Similarly, low level exposure to CO in air pollution (all levels below the World Health Organisation guideline of ~9ppm (World Health Organisation, 2000)) has been linked to a higher incidence of dementia (Chang et al., 2014), stroke (Hedblad et al., 2005; Maheswaran et al., 2005) and heart failure (Shah et al., 2013). Here, we used a range of target CO levels from 1ppm to 25ppm, capturing the lower, atmospheric level as well as guideline and higher exposures. Our high exposure level of 25ppm was aimed at half that of common commercial CO home alarm levels (50ppm).

As we observed a discrepancy between target and actual CO dose, we elected to present our findings according to the actual measured CO levels. The measured CO dose matched well with our moderate CO target (9.0ppm vs a 8ppm target), but less so with the highest and lowest levels of CO (18.2ppm vs a 25ppm target and 3.4ppm vs a 1ppm target, respectively), although it is important to note that actual measured CO revealed that embryos were exposed to low, moderate or high CO levels as intended. The reason for the discrepancy in target vs actual CO might be due to the gas delivery system, which utilised 20mL syringes to add the gas to the incubator boxes. For the lowest amount, the accuracy of such syringes might not be sufficient. For the highest amount, any inaccuracy in the delivery would be increased with an increase in dose. Future studies should endeavour to improve upon the delivery method to be able to address all levels of CO exposure.

### Dose dependency

A correlation analysis between measured CO (per individual batch) and heart wall thickness indicates that doses from 3.4ppm and up have an impact on heart wall thickness, and that this effect is linked to measured exposure dose. A significant positive correlation was seen between ventricular wall thickness and measured CO level across all experimental runs. A correlation was not seen between CO dose and heart weight. Heart weight appeared to drop with 3.4ppm and 9ppm, but rose again with 18.2ppm, which may explain this. This suggests that there is no linear relationship between CO dose and heart weight, but instead either a plateau is reached, or the weight loss might be offset by the increased ventricular wall thickness with dose. This adaptation could therefore explain the curvilinear relationship observed here.

### Comparison with other studies

This study is, to our knowledge, the first to examine low-level CO exposure effects on cardiac morphology in the chick embryo at a reasonably early stage of development (HH35). While ours is the first to look at very low levels of CO, other studies have looked at chick development during exposures to much higher doses of CO. Baker and Tumasonis identified a ‘critical level’ of CO concentration at 425 ppm for hatchability, at which point they observed no significant impact on egg weight, which concurs with our findings, nor congenital abnormalities, which were not formally assessed in our study (Baker & Tumasonis, 1972). The authors observed no effects when exposing eggs to <100 ppm, although no detailed investigation of cardiac development was done in the study (Baker & Tumasonis, 1972). Indeed, we observed no effect on viability for any of our exposure levels. In a subsequent study (Baker et al., 1973), the authors observed increases in hepatic oxidase enzymes in older embryos exposed to CO, which could represent an adaptation to hypoxia.

### Possible mechanisms

This study did not look at mechanisms, and so we cannot at present determine how CO causes the observed effect, e.g. whether the increase in wall thickness is due to hypertrophy or hyperplasia, and whether there is a change in extracellular matrix or inflammatory cells. Furthermore, CO is known to impact several important physiological pathways, meaning that there are several possible underlying mechanisms for the observed effects. For example, CO may impact the embryo through: hypoxia, oxidative stress, altered NO expression, endovascular inflammation, mitochondrial dysfunction and reactions with heme-containing proteins.

#### Hypoxia

CO overexposure is typically associated with hypoxia (Penney et al., 2010), and organs with a high oxygen requirement, such as the heart and the brain, may be especially susceptible to the effects of CO through this mechanism (National Research Council, 2010; Townsend & Maynard, 2002). Decreased oxygen delivery due to CO exposure can be compensated to some degree by e.g. increases in cardiac output and extraction of oxygen, but these mechanisms may be insufficient to fully restore oxygen delivery to the tissues. Development is a dynamic process, in which the embryo shows a high degree of plasticity. It is possible that an attempted compensatory increase in myocardial contractility in our CO-exposed embryos may be linked to the observed dose-dependent increase in ventricular wall thickness. Indeed, hypoxia at 10% O_2_ has been shown to cause hypertrophy in cultured neonatal rat cardiac myocytes, accompanied by an increase in hypoxia-inducible transcription factor-1 (specifically, HIF-1α) mRNA and protein levels (Chu et al., 2012). HIF-1α regulates the body’s cellular and developmental response to hypoxia. Studies have shown that CO can increase protein levels of HIF-1α (Choi et al., 2010; Rosenberger et al., 2007) as well as its isoform HIF-2α (Wiesener et al., 2003), promoting vascular endothelial growth factor (VEGF, an angiogenic factor) production (Choi et al., 2010). It must, however, be noted that the above studies used non-gaseous CO-releasing molecules (Choi et al., 2010) or gaseous CO at levels that were orders of magnitude higher (1000ppm) than those in the current study (Rosenberger et al., 2007; Wiesener et al., 2003).

As level of HIF isoforms were not assessed in the present study, we can only speculate as to their involvement in our findings, and we cannot yet with certainty determine to which extent hypoxia (and/or modulation of HIF-1α or HIF-2α) is the driver of pathophysiology development, particularly at the lowest levels of CO. The lowest levels of CO exposure in the current study were below common ambient air exposures in e.g. urban areas and may be too low to induce a strong enough hypoxic state to cause pathophysiology. Thus, we will include a discussion about other potential underlying mechanisms for the observed effect not related to hypoxia below.

#### Oxidative stress and NO expression

CO exposure may lead to oxidative stress through the effects of nitric oxide (NO). It is known that CO can increase inducible NO synthase (iNOS) expression, and iNOS expression may mediate myocardial damage (Rose et al, 2017). Cardiac dysfunction may be caused by inhibition of oxidative phosphorylation and CO binding to myoglobin (Rose et al, 2017). *In vitro* work in rats has shown that CO levels as low as 10.5ppm may significantly increase NO levels (Thom & Ischiropoulos, 1997), through the modulation of nitric oxide synthase (Thom et al., 2004). Work in the same model organism shows that NO can generate peroxynitrite, a potent free radical and oxidant (Ischiropoulos et al., 1996). This has been further described in studies on CO poisoning: after CO poisoning, NO-derived oxidants are needed for neutrophil adherence to the brain endothelium (Thom et al., 2001) and platelet-neutrophil aggregates and increased plasma myeloperoxidase, a biomarker of inflammation, is found in patients (Thom et al., 2006). It has been suggested that an immunological cascade involving oxidative species may be an underlying factor in delayed neurological sequelae after CO poisoning (Thom et al., 2004).

#### Heme-containing proteins and mitochondrial dysfunction

CO may react with heme-containing proteins other than Hb. Such targets include myoglobin and neuroglobin, NO synthase, NADPH oxidase, cytochrome c oxidase, cytochrome P450 and reactive oxygen species such as peroxidase (for an overview, see (Wu & Wang, 2005)). These heme-containing proteins are important for several physiological functions, from mitochondrial respiration to signal transduction. For example, a study in mice have shown that cytochrome c release is impaired in newborn mice exposed to levels of CO comparable with those in the present study, and that this is associated with neurodevelopmental impairment (Cheng et al., 2012). The toxicity of CO to mitochondrial function has been shown in humans (Alonso et al., 2003). CO may cause mitochondrial dysfunction through binding to cytochrome oxidase, and it may release NO and form peroxynitrite, thus causing further inactivation of mitochondrial enzymes and endothelial damage. It is possible that this could be amplified by further immunological responses. In a recent review, Almeida et al (Almeida et al., 2015) suggest that CO affects mitochondrial quality control machinery through reactive oxygen species signalling. Downstream effects of CO poisoning and the potential related mitochondrial dysfunction may also induce inflammatory signalling pathways (Rose et al., 2017) While some studies suggest that low doses of CO protects mitochondria from oxidative stress, the effect of CO on mitochondrial function is still poorly understood (Almeida et al., 2015).

While the above may play a role in the impact of CO on embryo development, further study is needed to identify which of these mechanisms are involved in pathophysiology development. Additionally, future work needs to take particular care to unpick the role of CO as a teratogen in its own right, or whether the downstream effects of hypoxia are the root cause

### Clinical relevance

In order to assess the impact of CO on health, it is important to know what dose produces which effect, and if this is uniform across the population. This is particularly important during pregnancy, as there is evidence to suggest that foetal tissues may be more vulnerable to hypoxia than maternal tissues. Indeed, CO levels that are not toxic to the mother may still be toxic to the developing foetus (Aubard & Magne, 2000; Farrow et al., 1990; Tomaszewski, 1999). There also remains the possibility that CO impact is not immediate, but rather that delayed sequelae may occur — as suggested in studies on the cognitive effects of CO poisoning (Choi et al., 2021; Jasper et al., 2005; Sönmez et al., 2018).

### Limitations

This study is the first to look at low-level CO impact on the heart in the developing chick embryo. While we observe an effect of CO on heart weight and ventricular wall thickness, the method of exposure means that CO levels did fluctuate throughout the 10-day exposure periods. It should be noted, however, that exposure levels never exceeded the target, and we are thus confident that the current findings truly reflect low-level exposure effects. To account for any possible effect on CO variation, we assessed the correlation between ventricular wall thickness and absolute (measured) CO levels. This second analysis showed significant correlation between wall thickness and CO exposures across all protocols. Future studies, however, should endeavour to minimise such variations in ambient gas composition. Furthermore, a repeat of the experimental protocol using the moderate CO exposure level showed the same increase in ventricular wall thickness compared to air control, suggesting that our findings are valid and repeatable.

Questions remain about the underlying cause of this effect as well as impact on other cardiac metrics. Future studies might wish to include assessment of effects on e.g. atrial wall thickness, lumen size and ventricular septum thickness, as well as interrogate the mechanism(s) underlying this impact.

### Conclusion

Low-level CO exposure *in ovo* for the first ten days of gestation impacts cardiac development in the chick embryo. Further studies are needed to fully elucidate the impact of low-level CO on cardiac development.

## ACKNOWLEDGEMENTS

We would like to thank Professor Marysia Placzek at the University of Sheffield School of Biosciences for her generous support of the experimental work and her valuable feedback on the manuscript. We would like to thank Dr Alex Fletcher (University of Sheffield) for his comments on the manuscript and support with planning the work. We are also very grateful to Oliver Smith Plumbing & Building (Sheffield) for their generous gift of a CO monitor for this work. Filipa Matias was supported by the Erasmus programme. Ian Groves was supported by the Engineering and Physical Sciences Research Council (PhD studentship). Mari Herigstad is funded by Kane International Ltd and the CO Research Trust.

## Notes

### Competing Interest Statement

The authors have declared no competing interest.

### Summary of Updates

Minor error in labelling of table/figures corrected.

